# A First-Order Design Requirement to Prevent Edema in Mechanical Counter-Pressure Space Suit Garments

**DOI:** 10.1101/633917

**Authors:** Christopher E. Carr, Loretta Treviño

**Author notes:** MIT, 77 Massachusetts Avenue, Room 54-418, Cambridge, MA 02139,. Senior CFD Engineer, TLG Aerospace, Seattle Washington. Received 2^nd^ place in Aerospace Medical Association Young Investigator Award in 2005 while an MIT Undergraduate for earlier draft of this manuscript.

## Abstract

**Introduction:** Mechanical counter-pressure (MCP) space suits may provide enhanced mobility relative to gas-pressure space suits. One challenge to realizing operational MCP suits is the potential for edema caused by spatial variations in the applied body-surface pressure (dP). We determined a first-order requirement for these variations.

**Methods:** Darcy’s law relates volume flux, of fluid from capillaries to the interstitial space, to transmural hydraulic and osmotic pressure differences. Albumin and fibrinogen levels determine, to first order, the capillary oncotic pressure (COP). We estimated dP, neglecting hydrostatic pressure differences, by equating the volume flux under MCP and under normal with the volume flux under abnormal variations in COP; then we compared these estimates to results from MCP garment studies.

**Results:** Normal COP varies from 20-32 mm Hg; with constant hydraulic conductivity, dP≈12 mm Hg. In nephrotic syndrome, COP may drop to 11 mm Hg, yielding dP≈15 mm Hg relative to mid-normal COP. Previous studies found dP_max_ =151 mm Hg (MCP glove; finger and hand dorsum relative to palm), dP_max_=51 mm Hg (MCP arm; finger, hand dorsum, and wrist relative to arm), and dP=52, 90 and 239 mm Hg (three MCP lower leg garments).

**Conclusions:** MCP garments with dP_max_≤12 mm Hg are unlikely to produce edema or restrict capillary blood flow; however, garments with dP_max_>12 mm Hg will not necessarily produce edema. For example, the hydrostatic pressure gradient at the feet in 1g can range from 70-90 mm Hg. Current garment prototypes do not meet our conservative design requirement.

## INTRODUCTION

In 1968-1971, Dr. Paul Webb demonstrated an alternative to gas pressure space suits, known as mechanical counter-pressure (MCP) [1,24]. While subjects wearing the MCP prototype Space Activity Suit (SAS) demonstrated excellent mobility relative to gas pressure space suits, subjects required extensive assistance in donning and doffing the SAS [1]. Altitude chamber tests of the SAS [1], like more recent investigations of MCP garments [3-4,20-21], were of quite limited duration (all <1 hour in final depressurized state in which the MCP garment was expected to provide a safe level of counter-pressure). Frank edema resulted during tests of the SAS [1] and during a 30-minute evaluation of an MCP glove [4], but not during a 5-minute evaluation of a different MCP glove [20]. The compatibility of MCP with longer-term fluid balance is unknown, as are the allowable variations in the body surface pressure, which can lead to edema in under-pressurized regions [1] or loss of blood flow and tissue ischemia in areas of over-pressurization [20].

In order to develop a practical space suit based on MCP, several hurdles must be surmounted: First, a practical donning and doffing solution must be identified. Second, the mean applied pressure must be approximately 200 mm Hg or greater in order to permit oxygen breathing pressures of the same order [16]. Third, the skin must not become overly irritated during extended contact with the MCP garment. Fourth, variations in body surface pressure should not produce significant tissue damage or discomfort in an acute or chronic setting, for example, with repeated use of an MCP space suit for extravehicular activity periods up to eight hours in duration.

In this paper, we develop a conservative design requirement for allowable variations in MCP garment body surface pressure and compare this design requirement to surface pressure data from past MCP garment studies.

## METHODS

To develop a conservative design requirement for the allowable variations in body surface pressure under conditions of chronic MCP garment use, we used Darcy’s law to construct a simple model of how variations in MCP body surface pressure may impact the microcirculation. We then collected data on edema formation under disease states, and used this data to estimate an equivalent variation in body surface pressure in the normal state. Finally, we computed the variation in body surface pressure for several MCP garments and compared these values to the normal and diseased states.

### Microcirculation Model

Darcy’s law, known as the Starling hypothesis when applied to fluid passage across the capillary endothelium [2], can be expressed as

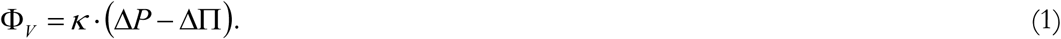

The hydraulic conductivity (filtration coefficient) of the capillary endothelium, *κ*, controls the volume flux of solvent (water and small molecules in the blood plasma), Φ_*V*_, driven by the net difference of the transmural hydraulic pressure difference, Δ*P*, and the transmural oncotic pressure difference, ΔΠ. In a normal individual, the total-body hydraulic conductivity is approximately 0.0061 ml · min_-1_ · (100 g tissue)_−1_ · (mm Hg)_−1_ [2]. As shown in Figure 1(A), we define Φ _*V*_ to be positive for flow out of the capillary into the interstitial space, and therefore define Δ*P* = *p*_*c*_ − *p*_*i*_ and ΔΠ = *α* (*π* _*c*_ − *π* _*i*_), where *p*_*c*_ is capillary hydraulic pressure, *p*_*i*_ is interstitial hydraulic pressure, *π* _*c*_ is capillary oncotic pressure, *π* _*i*_ is interstitial oncotic pressure, and *α* is the reflection coefficient. We set Δ*P* = 0 for *p*_*i*_ > *p*_*c*_ as a simple model of capillary collapse.

**Figure I.**
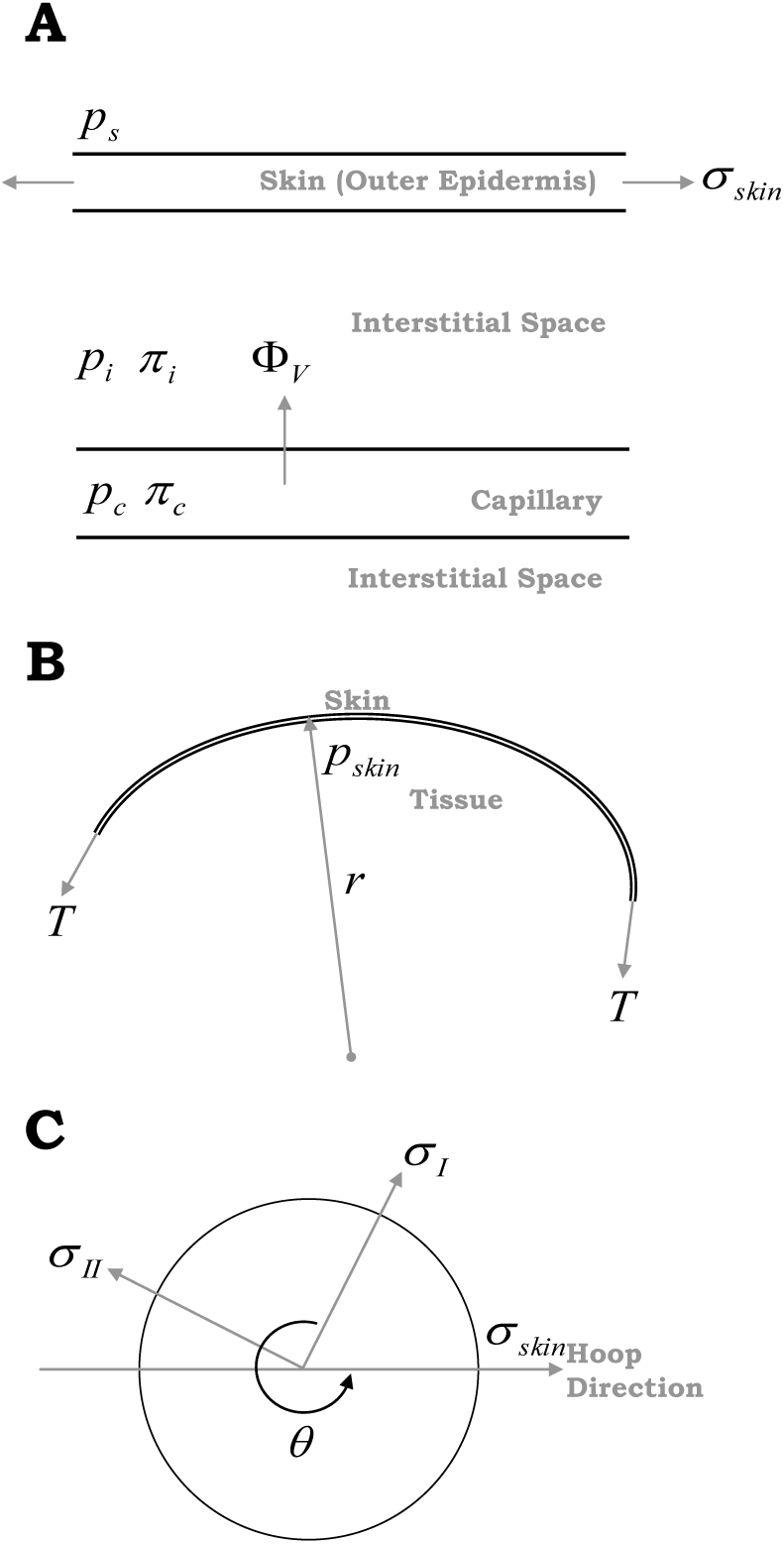
A. Physiological correspondence of variables used in one-dimensional microcirculation model. B. Modeling of a body segment as an irregular cylinder permits estimation of the contribution of skin tension to interstitial pressure. C. Transformation of stress for estimation of hoop direction skin stress. All variables are described in the text.

Capillary oncotic pressure, also called plasma colloid osmotic pressure, has been found to be correlated (*r* = 0.96, p < 0.001) with the sum of plasma Albumin and Fibrinogen concentrations and can be predicted as [9]:

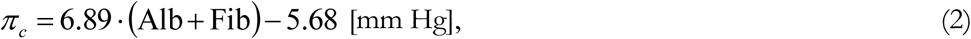

where Alb and Fib are the respective plasma concentrations of Albumin and Fibrinogen in g/dL. Fauschald et al. (1985) measured subcutaneous interstitial oncotic pressure to be 15.8±2.7 mm Hg in ten healthy humans [7]; we have taken *π*_*i*_ = 16 mm Hg except where otherwise indicated. We also took *α* = 1 because the reflection coefficient approaches unity for large proteins [2] such as Albumin and Fibrinogen.

Capillary hydraulic pressure can be estimated as *p*_*c*_ = *p*_*c*,0_ + *p*_*h*_. The first term, *p*_*c*,0_, typically varies from about 30 mm Hg to 10 mm Hg along the length of a capillary (the extrema depend strongly upon the resistance of pre-capillary and post-capillary vessels). We neglected hydrostatic pressure *p*_*h*_.

When no MCP garment is worn, subcutaneous interstitial hydraulic pressure, *p*_*i*_, is generally close to zero (relative to nominal atmospheric pressure), but skin tension elevates interstitial hydraulic pressure above zero in convex regions. Figure 1(B) illustrates the geometry used to estimate the contribution of skin tension to interstitial hydraulic pressure by treating a body segment as an irregular cylinder. Skin tension per unit length, *T*, can be estimated, using data from the literature, as *T* = *σ* _*skin*_ · *t*, where *t* is the skin thickness, and *σ* _*skin*_ is the in-vivo skin stress in the ‘hoop’ direction. The local skin radius of curvature, *r*, and tension per unit length, *T*, determine the hydraulic pressure applied by the skin to the interstitial space, resulting in an interstitial pressure of *p*_*i*_ = *T / r*.

Reihsner & Menzel (1996) reported in-vivo skin stress values based on 30 mm diameter skin samples taken from eight anatomical sites of six humans within 24 hours of death [19]. They reported the mean principle stresses *σ* _*I*_ and *σ* _*II*_, the maximum and minimum stresses, respectively. Using the standard transformation equations for stress [5] the ‘hoop’ stress can be estimated as:

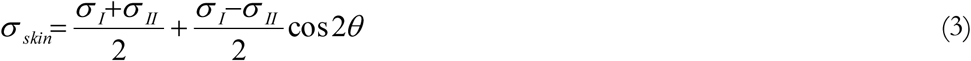

where *θ* is the angle of rotation, shown in Figure 1(C), required to rotate the physical coordinate axis of the principle strains so that the rotated axis of *σ* _*I*_ is aligned with the hoop direction.

Because *θ* is a strong function of body location, related to the Langer lines [14], we treated *θ* as a uniform random variable with range 0 ≤ *θ* < 2*π*. Because

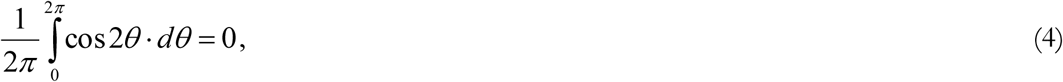

we approximated skin ‘hoop’ stress as

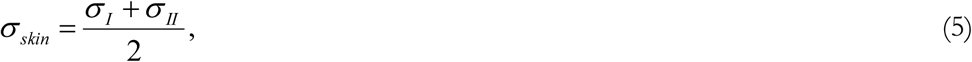

and used principle stress data from dorsal and volar thigh skin samples (of the available body locations sampled by Reihsner & Menzel (1996), the thigh best approximates a cylinder). Based on selection of thigh data we took *r* = 0.089 m by treating the thigh as a cylinder and using as the circumference the average of 50^th^ percentile male and female thigh circumference [15].

When an MCP garment is worn on an approximately cylindrical body segment, the subcutaneous interstitial hydraulic pressure can be estimated as:

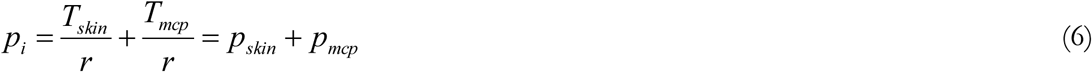

where *T*_*skin*_ is the skin tension per unit length, *T*_*mcp*_ is the MCP garment tension per unit length, *p*_*skin*_ = *T*_*skin*_ */ r* is the pressure applied by the skin, and *p*_*mcp*_ = *T*_*mcp*_ */ r* is the pressure applied by the MCP garment. In past MCP garment studies, investigators measured *p*_*mcp*_ using a force sensing array (Tekscan, South Boston, MA) placed between the skin and the MCP garment.

After estimating *p*_*skin*_, we used the one-dimensional microcirculation model to simulate a capillary of a normalized length of one, and computed Φ_*V*_ along the length of the capillary to illustrate, qualitatively and quantitatively, the effects of changes in oncotic pressure and the effects of spatial variations in pressure applied by MCP garments. In the two-dimensional model, oncotic pressures *π* _*c*_ and *π* _*i*_ are assumed constant along the length of the capillary, while *p*_*c*_ varies along the length of the capillary. Variables refer to capillary entry conditions except where noted. A bar (−) is used to indicate a mean quantity along the capillary; for example, 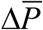 is the mean transmural hydraulic pressure difference along the capillary. The subscript _*net*_ indicates a quantity integrated along the capillary; for example, Φ_*V, net*_ is net volume flux.

### Hydraulic pressure equivalent of oncotic pressure changes

Because of the equivalence of oncotic pressure and hydraulic pressure, oncotic pressure changes in altered states provide some information about the allowable variation in MCP body surface pressure. Normal variations in oncotic pressure provide a conservative envelope of allowable variations, while oncotic pressures observed in edematous disease states provide evidence of the scale of variations that might produce edema in chronic MCP garment use. One can define:

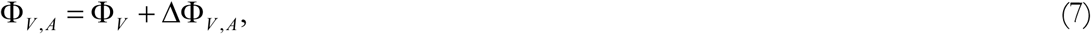

where Φ_*V, A*_ is the volume flux in state A, Φ_*V*_ is the volume flux in the normal state, and ΔΦ_*V, A*_ is the change in volume flux due to state A, relative to the normal state. By definition, one can write:

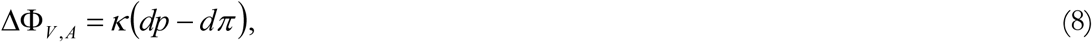

where *dp* is the change in transmural hydraulic pressure due to state A, and *dπ* is the change in transmural oncotic pressure due to state A. To derive equivalence between changes in oncotic pressure in altered states and equivalent pressure variations in MCP garments we set,

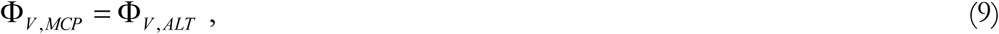

where Φ_*V, MCP*_ is the volume flux for a healthy person wearing an MCP garment, and Φ_*V, ALT*_ is the volume flux for a person in an altered (normal or disease) state not wearing an MCP garment. By definition, ΔΦ_*V, MCP*_ = ΔΦ_*V, ALT*_. If hydraulic conductivity is constant, an invalid assumption in some edematous disease states, and one assumes that ΔΦ_*V, MCP*_ = *κ* · *dp* and ΔΦ_*V, ALT*_ = −*κ* · *dπ*, then *dp*_*MCP*_ = −*dπ* _*ALT*_. In the general case (both *dπ* and *dp* are allowed to vary in the altered state), the equivalent *p*_*mcp*_ is given by 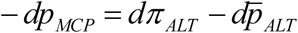.

We calculated *dπ* to estimate the allowable *p*_*mcp*_ from values of *π* _*c*_ and *π* _*i*_ available in the literature. Because synthesis of Albumin and Fibrinogen is known to co-vary [18], the normal *π* _*c*_ range was estimated using high normal (Fib = 0.4 g/dL, Alb = 5.0 g/dL) or low normal (Fib = 0.2 g/dL, Alb = 3.5 g/dL) values for both Fibrinogen and Albumin.

In all disease states examined except congestive heart failure we maintained the nominal capillary hydraulic pressure gradient of 30-10 mm Hg. In congestive heart failure, venous pressure is elevated substantially; because post-capillary resistance is much lower than pre-capillary resistance, changes in venous pressure have a much greater impact on capillary hydraulic pressure than changes in arterial pressure. To simulate this effect we reduced the capillary hydraulic pressure gradient for the congestive heart failure state by allowing *p*_*c*,0_ to vary from 30-20 mm Hg. Using the capillary model, we estimated Φ_*V, ALT*_ under each altered state.

### MCP Garment Evaluation

We evaluated pressure data from all known past MCP garment studies as of 2005 for which quantitative pressure data was published [3, 20-21]. Annis & Webb (1971) and Clapp (1983) did not report quantitative pressure data. From the available data, we computed the largest reported MCP garment pressure, *dP*_+_, the smallest reported MCP garment pressure, *dP*_−_, the maximum spatial variation in reported MCP garment pressure, *dP*_max_ = *dP*_+_ − *dP*_−_. Using the capillary model, we computed Φ_*V, MCP*_ applied pressures of *dP*_+_ and *dP*_−_. We report MCP garment pressure relative to atmospheric (breathing) pressure for the studies in which the MCP garment was worn within a partial vacuum chamber [3, 20-21]. Also analyzed was unpublished data from a study in which several MCP garments were tested on a non-compliant three-dimensional model of a human leg (custom built with the same leg geometry as the leg of the subject for whom the MCP garments had been designed) without a partial vacuum chamber; in this case we reported *p*_*mcp*_ relative to the spatial mean of the applied counter-pressure.

## RESULTS

Based on the normal range of Fibrinogen and Albumin, normal capillary oncotic pressure, *π* _*c*_, varies from approximately 20-32 mm Hg. Skin tension contributes on the order of 5 mm Hg to interstitial pressure in the thigh region (Table I).

**TABLE I.** SKIN TENSION CONTRIBUTION TO INTERSTITIAL PRESSURE.

| Sample Site | $\sigma_I$<br>kPa | $\sigma_{II}$<br>kPa | $\sigma_{skin}$<br>kPa | $t$<br>mm | $T_{skin}$<br>N/m | $r$<br>Mm | $p_{skin}$ | |
| --- | --- | --- | --- | --- | --- | --- | --- | --- |
|  |  |  |  |  |  |  | kPa | mm Hg |
| Dorsal Thigh | 44.4 | 28.5 | 36.5 | 1.6 | 58.3 | 8.9 | 0.66 | 4.9 |
| Volar Thigh | 55.6 | 20.7 | 38.1 | 1.5 | 57.2 | 8.9 | 0.64 | 4.8 |

Reihnsner & Menzel (1996) reported skin principle stresses ( $\sigma_I, \sigma_{II}$ ) in newtons [19]; the stresses reported here were computed using the 30 mm sample diameter and skin thickness $t$ . Other variables are described in the text.

Figure II shows the results of the microcirculation model under six representative conditions, including low normal and high normal *π* _*c*_, positive and negative variations in MCP pressure, and a combination of negative MCP pressure and elevated capillary oncotic pressure. The resulting Φ_*V, net*_ is a linear function of the capillary inlet conditions except under conditions for which Δ*P* reaches zero and capillary collapse occurs, as in Figure II(F).

**Figure II.**
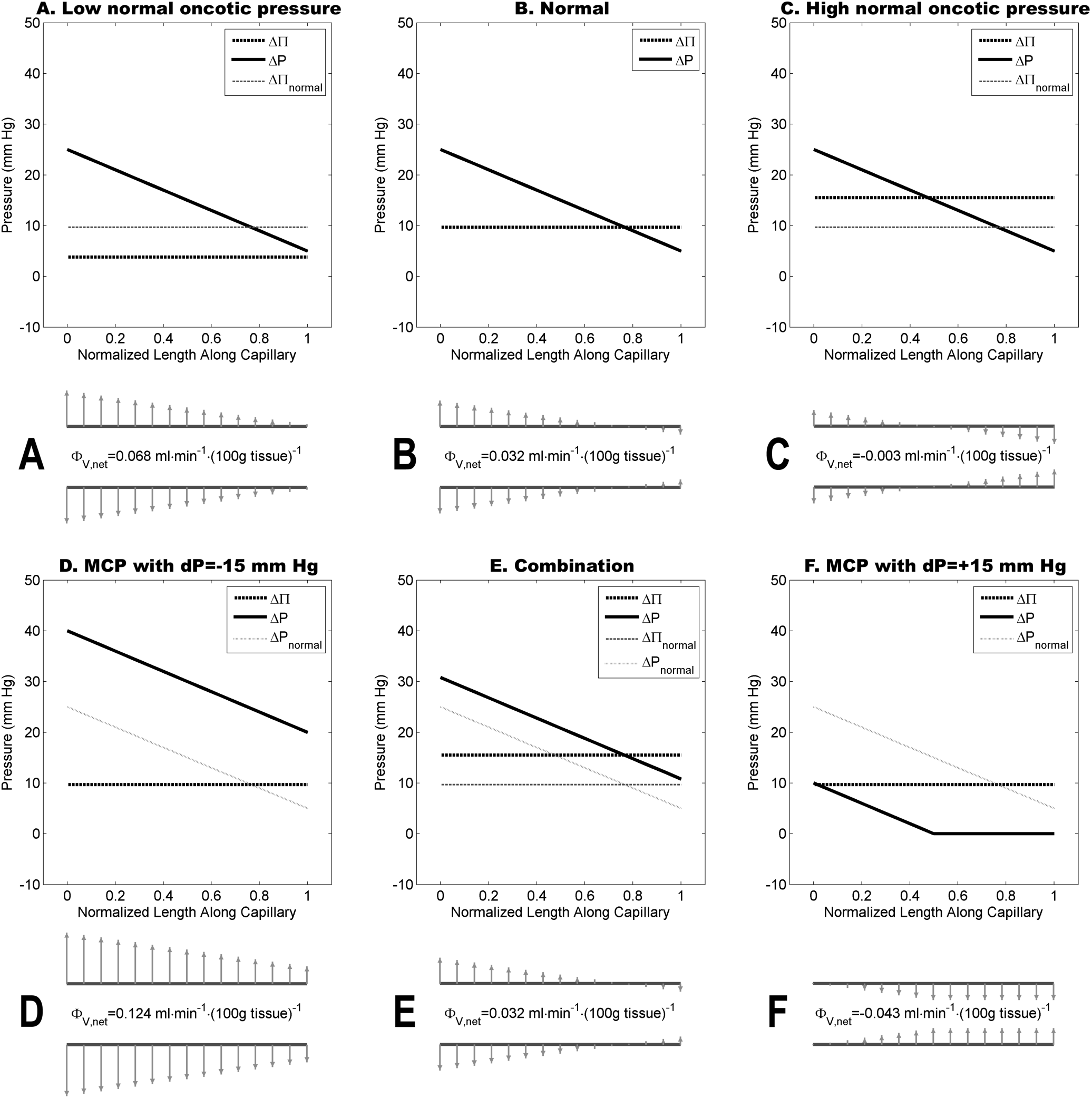
Microcirculation model applied to a capillary of a normalized length of one under six representative conditions: transmural hydraulic pressure Δ*P* and transmural oncotic pressure ΔΠ, the pressure terms of Equation 1, are plotted for each condition (A-F) along with a representative capillary below each plot. Grey arrows indicate the relative magnitude of the local volume flux Φ_*V*_ at points along the length of the capillary. In condition E (Combination), *p*_*mcp*_ = −5.8 and *dπ* = 5.8.

Hydraulic pressure equivalents to edematous disease states, excluding congestive heart failure, ranged from about −8 to −15 mm Hg (Table II). The *dP*_−_ values for MCP garments are, on average, a factor of 3-6 times greater in magnitude than this range (Table III). The *dP*_+_ values for MCP garments are, on average, a factor of approximately 13 times greater in magnitude than the equivalent *dP*_+_ resulting from normal variations in capillary oncotic pressure. For the MCP Glove, *dP*_max_ represents applied pressure at the finger and hand dorsum (not significantly different) relative to the palm. For the MCP Arm, *dP*_max_ represents applied pressure at the finger, hand dorsum, and wrist (not significantly different) relative to the arm; no palm pressure was reported for the MCP Arm. MCP Lower Leg conditions 1 and 2 involve the identical MCP garment; placing the garment on a three-dimensional model of a human leg (condition 2) resulted in a 4.4-fold increase in *dP*_max_ relative to placing the garment on a compliant human leg (condition 1).

**TABLE 2.** ONCOTIC PRESSURE CHANGES IN ALTERED STATES.

| Condition | Albumin<br>g/dL | Fibrinogen<br>g/dL | $\pi_c$<br>mm Hg | $\pi_i$<br>mm Hg | $d\pi$<br>mm Hg | Reference |
| --- | --- | --- | --- | --- | --- | --- |
| Low normal $\pi_c$ | 3.50 | 0.20 | 19.8 | 16 | -5.8 | (Not applicable) |
| Normal | 4.25 | 0.30 | 25.7 | 16 | 0.0 | (Not applicable) |
| High normal $\pi_c$ | 5.00 | 0.40 | 31.5 | 16 | +5.8 | (Not applicable) |
| Kidney Failure | 2.50 | 0.30* | 13.6 | 16 | -12.1 | Post & Rose (2004) |
| Nephrotic Syndrome |  |  | 10.9 | 16 | -14.8 | Koomans et al. (1986) |
| Hepatic Cirrhosis - Controls | 3.10 | 0.30* | 17.7 | 16 | -7.9 | Güven et al. (1993) |
| Hepatic Cirrhosis - Survivors | 3.40 | 0.30* | 19.8 | 16 | -5.9 | Güven et al. (1993) |
| Hepatic Cirrhosis - Non-survivors | 2.40 | 0.30* | 12.9 | 16 | -12.7 | Güven et al. (1993) |
| Congestive Heart Failure | 3.57 |  | 22.3 | 1.0 | +11.6<sup>†</sup> | Kwan et al. (1990) |

\* Normal Fibrinogen levels assumed in estimation of $\pi_c$ . <sup>†</sup> In this state, $d\bar{p} = 5.0$ mm Hg due to elevated venous pressure, yielding an equivalent $p_{mcp}$ of 6.6 mm Hg; $d\bar{p} = 0$ for all other disease states.

**TABLE 3.** MCP GARMENT SPATIAL PRESSURE VARIATIONS.

| Garment or Condition | Description | $dP_+$<br>mm Hg | $dP_-$<br>mm Hg | $dP_{\max}$<br>mm Hg | Reference |
| --- | --- | --- | --- | --- | --- |
| Normal | Normal range of $\pi_c$ | 5.8 | -5.8 | 12 | (Not applicable) |
| MCP Glove | Locations: F, D, P* | 74 | -78 | 153 | Tanaka et al. (2002) |
| MCP Arm | Locations: F, W, D, FA, UA* | 78 | 23 | 55 | Tanaka et al. (2003) |
| MCP Lower Leg 1 | 1-Channel Garment<sup>†</sup> | 40 | -14 | 54 | Bethke et al. (2004) |
| MCP Lower Leg 2<sup>‡</sup> | 1-Channel Garment<sup>†</sup> | 147 | -92 | 239 | Unpublished data |
| MCP Lower Leg 3<sup>‡</sup> | Foam-Bladder Garment<sup>†</sup> | 33 | -20 | 52 | Unpublished data |
| MCP Lower Leg 4<sup>‡</sup> | Multi-Channel Garment<sup>†</sup> | 67 | -23 | 90 | Unpublished data |

\* All measurement locations reported by authors are given: F=top of finger, D=dorsum of hand, P=palm of hand, W=wrist, FA=forearm, UA=upper arm. Calculation based on mean pressure at reported locations under -150 mm Hg chamber pressure. <sup>†</sup> Applied pressure was measured for the MCP Lower Leg garments using a pressure measurement array (Tekscan, South Boston, MA); calculation based on pressure at a given height above the ankle averaged over circumference of lower leg. <sup>‡</sup> Because a partial-vacuum chamber was not used, these MCP lower leg garment pressure measurements were made on a three-dimensional model of a human leg instead of an actual human leg.

Net volume flux, Φ_*V, net*_, is greater in magnitude than in the normal state in all disease states except the condition of Hepatic Cirrhosis – Survivors (Fig. III). If the volume flux extrema in the normal state are taken to be ± 1*σ*, then the average volume flux in the disease states, excluding congestive heart failure, is 1.82*σ*, significantly greater than normal volume flux (*p* = 0.001). The volume flux extrema for MCP garments are entirely outside the normal range. For *dP*_+_ > 25 mm Hg, Φ _*V, net*_ reaches a negative extremum of − 0.059 ml · min^-1^ · (100 g tissue)^−1^ due to total capillary collapse in the model; all MCP garment conditions reached this negative extremum.

**Figure III.**
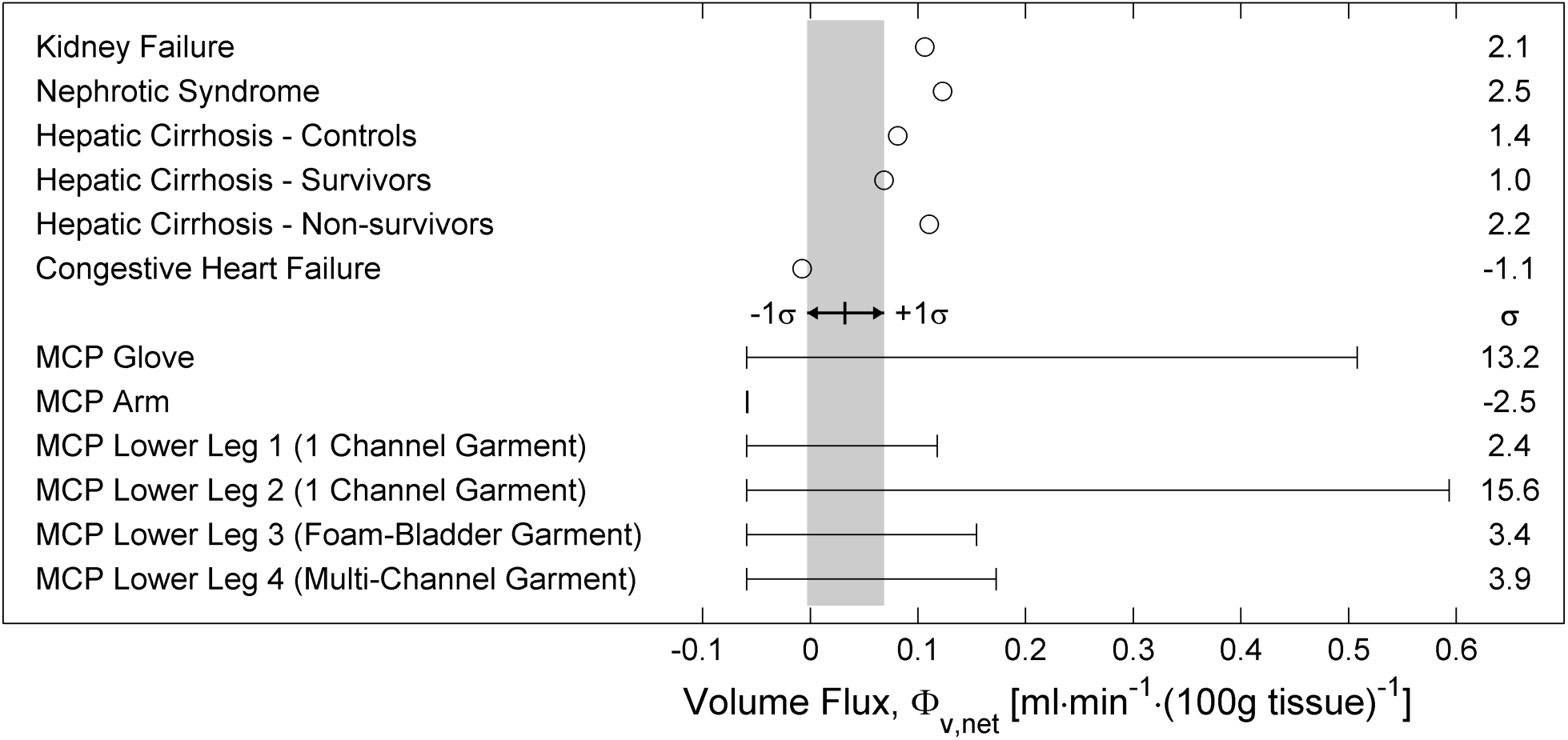
Volume flux computed using the capillary model, for disease states and for variations in body surface pressure due to the prototype MCP garments. For each MCP garment condition the left-most vertical line indicates the volume flux at the reported *dP*_+_, while the right-most vertical line indicates the volume flux at the reported *dP*_−_. The horizontal limits of the grey bar indicate the computed limits of volume flux (taken to be ± 1*σ*) due to normal variations in capillary oncotic pressure. Numbers at the right of the figure give the positive extreme of volume flux in terms of *σ*.

## DISCUSSION

The point of zero volume flux in the microcirculation model depends upon many parameters that vary spatially in the human body, and the actual balance between filtration and absorption is affected by many factors other than those included in the model. While healthy humans in normal conditions can tolerate large hydrostatic pressures of 70-90 mm Hg at the ankle during hours of sustained standing, without significant muscle activity edema can result. In comparison, 10° head-down tilt produces progressive facial edema, detectable after one hour [6].

The capillary model has at least two major deficiencies: the model is entirely passive, and the model of flow limitation based on capillary closure is a gross approximation. For example, the lack of feedback regulation causes the model to over-predict changes in volume flux due to changes in transmural hydraulic pressure. Normally, an increase in transmural hydraulic pressure results in an elevation of pre-capillary resistance, whereas a decrease in transmural hydraulic pressure results in a decreases in pre-capillary resistance [2]. Capillary closure would also limit flow, thereby affecting the pressure profile along the entire capillary length; our model of a partially collapsed capillary can be thought of as representing a collection of capillaries, with cross-connections enabling flow (up to a point) even with partial capillary collapse and flow limitation.

While one should interpret quantitative results of the model carefully, the model does provide plausible results, useful to understand the potential impact of variations in MCP garment pressure on the microcirculation. For example, in the normal condition volume flux is small and positive, consistent with removal of fluid from the interstitial space by the lymphatic system. The normal range of capillary oncotic pressure changes result in reasonable volume fluxes, with the high normal oncotic pressure condition representing a low rate of dehydration of the interstitial space (net absorption). While our rough calculation demonstrates the possibility that skin tension has a non-negligible impact on interstitial hydraulic pressure, the impact of skin tension on hydraulic pressure is likely to vary substantially as a function of body location, and deserves further study.

In edematous disease states other than congestive heart failure, the increased volume fluxes predicted by the capillary model, relative to the normal state, are consistent with expectations. In addition, moderately reduced capillary oncotic pressures were associated with worse outcomes. For example, in the cited study of hepatic cirrhosis [10], non-survivors had a *dπ* twice as negative as did survivors, who had low-normal capillary oncotic pressures. The state of congestive heart failure is qualitatively different than the other disease states, and we did not attempt to model all factors that affect fluid balance in congestive heart failure; in congestive heart failure, the low interstitial oncotic pressure acts to protect against the formation of edema [13], with the result that our model predicts a slightly negative volume flux in this state.

### MCP Garments

Because the MCP pressure data used in this analysis were all means of pressure measurements, actual spatial variability in the original pressure measurements is likely to be greater than the spatial differences reported here. However, the original pressure measurements were limited in quality by sensor noise, particularly the sensitivity of the Tekscan sensor outputs to bending or bunching of the sensors between the body surface and the MCP garment; this sensor noise may artificially increase the measured variability in the source data. The large increase in *dP*_max_ due to measurement of the 1-Channel MCP garment on the three-dimensional leg model versus a human leg illustrates the limitations of evaluating garments on a non-compliant model. This model may elevate the reported spatial variations by not deforming elastically, and it may also contribute to the reported spatial variations through additional bending of the sensors along its surface.

With these limitations in mind, the large relative magnitude of the spatial variations of the MCP garments, relative to the disease-state equivalent MCP garment pressures, suggests that similar MCP garments might have a significant impact on the microcirculation under conditions of chronic wear.

In the short-run, some of the reported negative variations in MCP garment pressures do not represent an extreme risk: subjects have tolerated up to four hours of −30 mm Hg of lower body negative pressure with initially increased but no net transcapillary fluid transport [23], suggesting that adequate compensatory mechanisms exist to cope with large body surface areas of moderately large negative deviations in MCP garment pressure for periods on the order of hours. However, in areas where MCP garment pressure is too low, such as flat areas like the palm of the hand that are difficult to pressurize, edema can result even over short periods. The occurrence of edema in a 30-minute duration MCP glove evaluation [4] and lack of edema in a five-minute duration MCP glove evaluation [20] may be due to differences in glove design, but more likely results from the difference in duration at the lowest chamber pressure. While the dense connective tissue of the palm may make it more resistant to edema formation [20], the variable resistance of body tissues to edema formation secondary to under-pressurization cannot be determined without longer duration tests of up-to-date MCP garments.

MCP garments may reduce blood flow to areas under high applied pressure, but at what applied pressure might this occur and how significant is the reduction in blood flow? Vollmar et al. (1999) applied external pressure to skin folds of Syrian golden hamsters as a model of compartment syndrome and determined the applied pressure required for blood flow to cease in different elements of the microcirculation using intravital fluorescence microscopy [22]. They found that blood flow ceased in 50% of the muscle capillaries at an external pressure of 12 mm Hg (our capillary model has a 50% collapse point at 15 mm Hg). Arteriolar flow cessation was a function of vessel size, with flow ceasing in vessels of diameter < 20µm at 26 mm Hg; although flow had ceased, arteriolar vessels did not collapse even at external pressures of 70 mm Hg. Blood flow ceased in venules between 27-33 mm Hg. In a separate study of nail capillary beds in humans, zero flow in the capillary beds was first observed at external pressures of 40 mm Hg during body cooling, 13 mm Hg during body heating, 28 mm Hg during regulated vasoconstriction, and 12 mm Hg during regulated vasodilatation [8].

If the maximum spatial variation of MCP garment pressure is limited to less than 12 mm Hg, the ability to adjust the breathing pressure provides a degree of freedom that can be used to ensure that the MCP garment provides no greater driving force to modify capillary fluid balance than do normal variations in capillary oncotic pressure. MCP garments meeting this criterion should not produce edema and would be unlikely to adversely impact capillary blood-flow, even with chronic usage. Garment prototypes as of this analysis (∼2005) fail to meet this design requirement by a factor of 4-20; with chronic usage it is possible, although not certain, that these garments could produce significant edema or impair capillary blood-flow and cause tissue ischemia. The 12 mm Hg maximum spatial variation criterion is clearly a restrictive design requirement, and we do not even venture to suggest that such a garment would be ideal. Garments using graduated compression are effective in reducing the edema associated with standing [12]; the trade-off between gradient versus non-gradient MCP garments has yet to be decisively examined.

